# Wake consolidates event memory, sleep binds it with context

**DOI:** 10.1101/2025.11.12.687983

**Authors:** Xiu Miao, Jessica Dahmen, Amelie Sophie Thoma, Anna-Lena Weber, Florian Waszak, Jan Born, Karsten Rauss

**Affiliations:** Institute of Medical Psychology and Behavioral Neurobiology, University of Tübingen, Germany; Graduate Training Centre of Neuroscience/IMPRS for Cognitive & Systems Neuroscience, University of Tübingen, Germany; Université Paris Cité, Integrative Neuroscience and Cognition Center, UMR 8002, Centre National de la Recherche Scientifique, Paris 75006, France; Werner Reichardt Centre for Integrative Neuroscience, University of Tübingen, Germany; German Center for Diabetes Research (DZD), Institute for Diabetes Research & Metabolic Diseases of the Helmholtz Center Munich at the University of Tübingen (IDM), Germany; German Center for Mental Health (DZPG) Tübingen, Germany

**Keywords:** memory consolidation, context, sleep spindles

## Abstract

Sleep is thought to be superior to wakefulness in consolidating memory, due to a hippocampus-driven process enhancing the contextual integration of events. We compared recall of associative memory in different contexts, and found that memory is also consolidated during wakefulness, although in a qualitatively distinct way. Healthy participants performed a cued stimulus-response (S-R) learning task requiring speeded object classification. Learning was followed by 2 hours of sleep or wakefulness. Reaction times revealed stronger S-R memory after sleep than wakefulness with retrieval in the same context, and this benefit was linked to increased sleep spindle activity. In contrast, S-R memory was stronger after wakefulness than sleep when tested in a different context. This wake-dependent enhancement had a fast onset and was reversed by delayed sleep. We conclude that, unlike sleep strengthening the binding of an event with its context, wakefulness stabilizes event representations itself, i.e., not bound to a specific context.

## Introduction

Sleep supports memory consolidation, i.e., the conversion of newly encoded memory traces into more stable forms that are easier to retrieve (Dudai et al., 2015). The consolidation of newly encoded experience during sleep is thought to be achieved in an active systems consolidation process that is based on the repeated neuronal replay of newly formed representations (Brodt et al., 2023; Diekelmann & Born, 2010; Lutz et al., 2025). This repeated replay leads to a gradual transformation of the originally encoded episodic memory representation, such that it more strongly involves neocortical networks. Systems consolidation and transformation of episodic memory representations during sleep crucially depend on thalamo-cortical spindles that occur during non-rapid eye movement (NonREM) sleep, precisely timed to the occurrence of ripples in the hippocampus, and to slow oscillations in the neocortex (Bergmann et al., 2012; Latchoumane et al., 2017), possibly facilitating the hippocampo-to-neocortical transfer of memory information.

Remarkably, recent findings have provided hints that memory consolidation can occur also in the wake state, although in a different way (Wamsley, 2019, 2022). In rats, memory for objects was even stronger after a post-encoding wake period than after a post-encoding sleep period, but only when retrieval was tested in a context different from that during encoding (Sawangjit et al., 2022). These findings suggest that whereas hippocampus-dependent consolidation during sleep strengthens the integration of items in memory with contextual information, consolidation in the wake state may enhance context-independent representations (Mednick et al., 2011; Sawangjit et al., 2018; Schapiro et al., 2019; Yonelinas et al., 2019). Indeed, these findings in rodents point to a central role of the processing of context information that may make wake consolidation superior to that during sleep in strengthening context-independent representations of an event (Maren, 2011).

Here, we aimed to show that in humans, like in rats (Sawangjit et al., 2022), consolidation during a post-encoding wake period surpasses consolidation during an equivalent post-encoding sleep period in strengthening context-independent memory of an event. As event we used a cued stimulus-response (S-R) task in which participants learned to classify visually presented objects as fast as possible with a preceding cue indicating (i) whether the object should be classified according to its size or its mechanical properties and (ii) whether to respond with a left or right button press (Fig. 1A, Moutsopoulou et al., 2018). A foregoing study proved that post-encoding sleep robustly strengthens the learned S-R associations on this task, as indicated by enhanced “switch costs” at the retrieval test, i.e., a distinctly slowed reaction time whenever an object is presented with a cue asking for a classification or response mapping different from that during the learning session (Fig. 1B; Miao et al., 2023). In order to test the context-dependency of these memories after offline sleep and wake intervals, we employed two different contexts, with the participants performing the task either in a virtual reality (VR) or real environment (RE), both of which were also associated with different experimenters and labs (Fig. 1C). As hypothesized, S-R memories were improved after post-encoding sleep when participants were tested in the same context as during encoding. Intriguingly, the opposite pattern was observed following wake consolidation: i.e., participants showed higher memory performance when tested in a different context, contradicting a long history of research on context-dependent memory (Godden & Baddeley, 1975; Smith, 1979, 1982; Smith et al., 1978).

**Figure 1.**
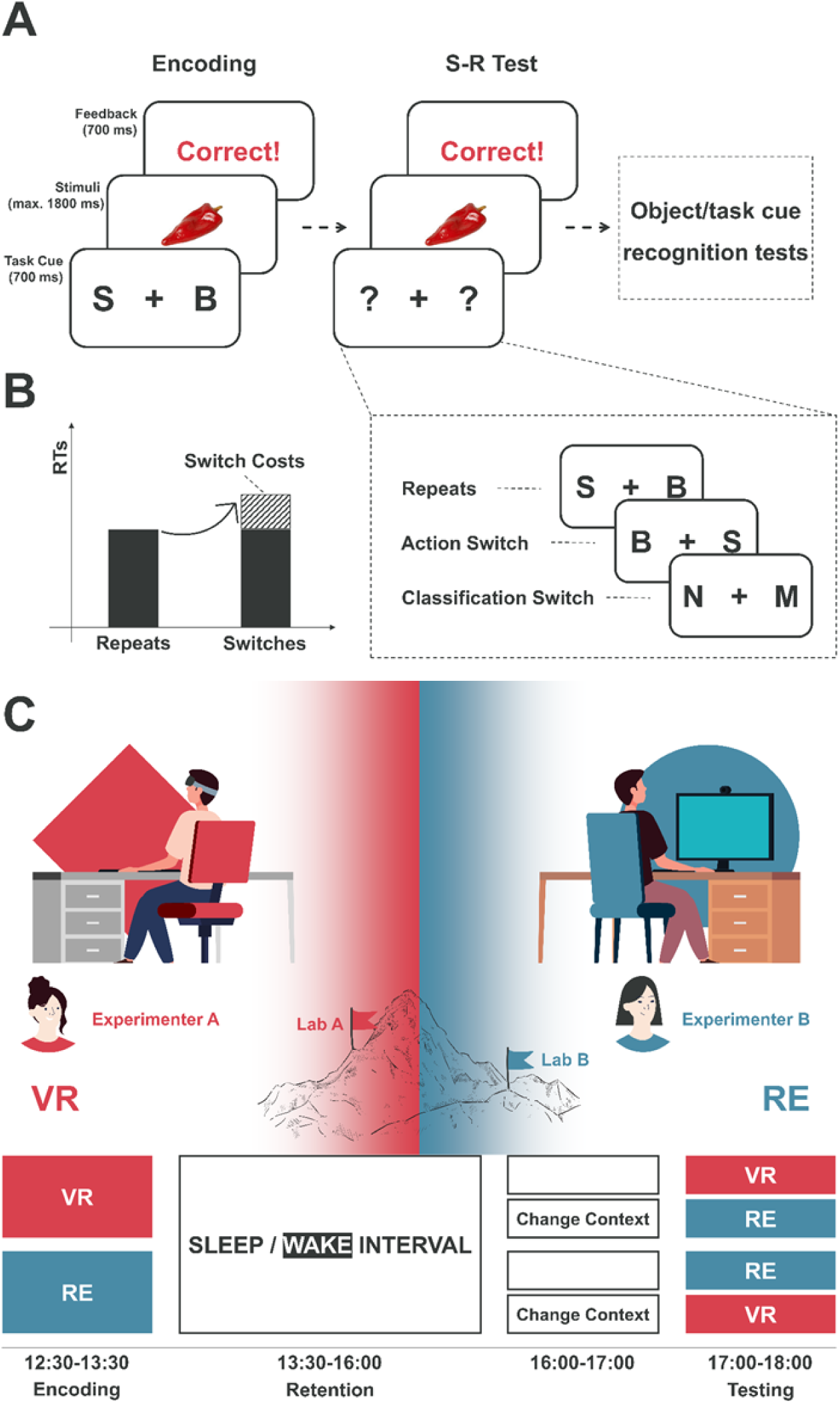
Tasks and procedures. **(A)** During encoding, each trial started with a cue indicating the task to be performed (i.e., to classify an object based on size or mechanical properties) and the response mapping, followed by presentation of an object. S (smaller) and B (bigger) refer to classification of the object’s size; M (mechanical) and N (non-mechanical) to classification of the object’s mechanical properties. For instance, the S displayed on the left tells the participant to press the left key when the object is smaller than a basketball, and to press the right key when it is bigger. Trials ended after a response was given or 1800 ms had elapsed with accuracy feedback (task instructions are translated from German). During testing, each object was associated with either the same task parameters as during encoding (repeat condition) or required a different motor response (action switch) or a different classification (classification switch). After S-R testing, we tested object recognition (by presenting both encoded and new objects, asking participants to make old/new judgments) and recognition of task cues associated with each object at encoding (by asking participants which of two task cues had been presented with the object during encoding). **(B)** Memory of S-R associations was measured in terms of switch costs, as indicated by the increase in RTs for switch trials (hatched) compared to repeat trials. **(C)** Participants either wore VR headsets to engage in a virtual space and completed tasks by viewing a virtual screen (VR, in red), or sat in front of a standard physical screen to complete the same tasks (RE, in blue). These settings were implemented in distinct labs located at different university campuses and by two different experimenters. After either taking a nap or staying awake during the retention interval, participants left and returned later to the same (Sleep: n = 17; Wake: n = 16) or a different (Sleep: n = 16; Wake: n = 17) context to be tested. The order of context manipulations was counterbalanced.

## Materials and Methods

### Participants

A total of 126 participants between 18 and 36 years old (mean age 23.15 ± 3.52 years, 27 men) took part in the experiments. None reported a history or presence of neurological or psychiatric disorders, drug abuse, or sleep problems. All participants maintained regular sleep/wake cycles and did not work on night shifts during the 4 weeks preceding the experiment. Participants were instructed not to consume alcohol or caffeine on the day of the experiment. Participants in the main experiment (including an experimental nap) indicated that they were able to nap during pre-screening. Two participants were excluded as they were not able to fall asleep during the nap period. Sample sizes were calculated using G*Power (Faul et al., 2009) based on previous studies with similar protocols (Miao et al., 2023; Moutsopoulou et al., 2018), to achieve 80% power at a 5% *⍺*-level. All participants were naive to the study paradigm and provided written informed consent before the experiment. The study was approved by the Ethics Committee of the Medical Faculty at the University of Tübingen and conducted in accordance with the principles of the Declaration of Helsinki (World Medical Association, 2013).

### Design and procedure – main experiment

The procedure for the main experiment was as follows: During a first encoding session, the participants performed on a stimulus-response (S-R) learning task. In the test session 3 hours later, (i) memory for S-R associations and (ii) recognition memory for the objects and (iii) the task cues associated with the objects in the encoding session was tested. A between-groups design was adopted with the participants either napping or staying awake during a 2-hour post-encoding interval and being tested in either the same context as during learning or in a different context, resulting in 4 groups (cf. Figure 1): sleep-same context (n = 17, 7 men); sleep-different context (n = 16, 1 man); wake-same context (n = 16, all women); and wake-different context (n = 17, 3 men).

Participants arrived at the lab at 12:30 h, received information and instructions, and signed the informed consent form. If the encoding context was the virtual reality (VR) setting, participants were first equipped with VR headsets and entered the virtual space. They were required to complete two adaptation tasks to acclimate to the virtual space, i.e., one minute of free movement using the controller to familiarize themselves with the surroundings, followed by watching a 9-min documentary displayed on the virtual screen to help them adapt to the state they would need to maintain during the experiment.

The encoding session took place in a quiet room, with the participants in the real environmental (RE) context seated ∼50 cm away from a standard monitor, and in the VR context seated in front of a table wearing VR headsets. After receiving task instructions, participants completed 16 practice trials on the S-R task to become familiar with the task. The stimuli used for practice were not presented in the main experiment. During subsequent encoding, each of 144 stimuli was presented twice in random order, resulting in a total of 288 trials (see task description below). Accuracy feedback was provided after each trial. Trials were divided into three blocks of equal duration, with breaks of 30 s between blocks. At the end of each block, average accuracy feedback was provided to further enhance motivation. Participants were free to initiate the next block after each break, allowing for longer breaks if needed. The encoding session lasted ∼30 min.

Following the encoding session, the Sleep groups were equipped with electrodes for polysomnographic recordings and allowed to sleep in the lab for two hours, while the Wake groups engaged in a series of standard activities that involved alternating between watching a 45-minute documentary and taking a 15-minute walk. The Sleep groups were awakened around 16:00 h. Afterward, both groups were instructed to leave the lab and return for the testing session one hour later, which ensured dissipation of sleep inertia (Achermann et al., 1996; Hilditch & McHill, 2019; Occhionero et al., 2021). They were instructed to avoid stressful physical or mental activities in this hour. Participants tested in a different context traveled from one lab to the other during this time.

Before the test session, participants in the VR setting completed the same VR adaptation tasks. Then, each of the encoded 144 stimuli was presented once, with either the same task cue as during the encoding session or a different task cue, resulting in either a repeat trial or a switch trial (see task description below). All trials were equally divided into three blocks, with breaks of 30 s between blocks, and participants received accuracy feedback after each block and could decide when to proceed to the next block, as during encoding. Following the S-R memory test, recognition memory for the objects and their associated task cues was tested. Testing of object recognition (288 trials) and task cue recognition (144 trials) was also divided into three blocks each, with breaks of at least 30 s between blocks. Accuracy feedback was only provided here on the object recognition task. In addition, a vigilance test (Diekelmann et al., 2013) and the Stanford Sleepiness Scale (SSS) (Hoddes et al., 1972) were applied at the start of the test sessions. The SSS revealed that the Wake group was subjectively more fatigue than the Sleep group prior to the test (*t* = −6.55, *p* < 0.001). Reaction times in the vigilance test were comparable between groups (*t* = −1.11, *p* = 0.271). No significant differences were found in the SSS and the vigilance test between context conditions (all *p* > 0.257).

### Supplementary experiments

Based on findings of the main experiment indicating an unexpected enhancement in S-R memory for the Wake group when tested in a different context, we performed two supplementary experiments that aimed to explore the temporal dynamics of this wake-dependent memory enhancement. In the first of these experiments, participants were tested closely (∼ 20 min) after encoding (Immediate-Wake groups) either in the same context (n = 20, 6 men) or a different context as during encoding. (n = 22, 4 men). The ∼20-min retention interval was owed to the fact that, for testing in the different context, participants had to travel to the other lab.

The second supplementary experiment aimed at testing the persistence of the wake-dependent increase in S-R memory (Delayed-Sleep group, n= 16, 6 men. Here, the encoding phase took place at 12:00 h, followed by a 12-hour wake-interval. The participants then slept at home, stayed there, and returned to the respective other lab for testing in the different context at 12:00 h, i.e., 24 hours after encoding. In the 12-hour wake period, the participants performed the same standard activities as described for the main experiments and then went home. Sleep duration was recorded using the MotionWatch (CamNtech Ltd., UK). In addition, participants were asked to complete an activity diary to document their specific activities while they were out of the lab.

The order of context manipulation was counterbalanced for both supplementary experiments. All other procedures in these supplementary experiments were as described for the main experiments. Reaction times in the vigilance test were comparable between all three supplementary groups (*F*(2, 55) = 0.27, *p* = 0.765).

### Tasks

We employed a cued S-R learning task, modified from a previous protocol (Fig. 1A, Miao et al., 2023). The tasks described below, including the instructions, were administered in German to native speakers, with only a few participants receiving an English version. For ease of understanding, the English version is presented below as an example. On this task, a total of 144 pictures of everyday objects were centrally displayed in color against a white background. Participants were asked to classify each stimulus based on either its real-life size or its mechanical properties, i.e., they had to decide whether the object shown was smaller or larger than a basketball, or whether it was a mechanical or non-mechanical object. For example, an avocado would be smaller than a basketball and non-mechanical object, while a motorcycle would be larger than a basketball and mechanical. Each stimulus was preceded by a cue consisting of two letters presented to the left and right of a central fixation cross. This cue indicated both the classification task to be performed on the next object, and the response side to be used. Thus, the letters ‘M’ on the left and ‘N’ on the right signaled a mechanical versus non-mechanical classification, where mechanical objects required a left-hand key press and non-mechanical objects required a right-hand response. Similarly, the letters ‘S’ (presented on the left) and ‘B’ (presented on the right) indicated a size classification task, with ‘smaller’ responses mapped to the left hand and ‘bigger’ responses to the right hand. The ‘A’ and ‘L’ keys on a standard keyboard were designated for left-hand and right-hand responses, respectively. Each trial began with the task cue—either ‘M/N’ (or ‘N/M’) or ‘S/B’ (or ‘B/S’)—which was presented for 700 ms, immediately followed by the central presentation of the object. The object remained on the screen for a maximum of 1800 ms or until a response was made. After each trial, participants received performance feedback (‘correct’, ‘wrong’ or ‘no response’) for 700 ms. Additionally, average accuracy was displayed at the end of each block of trials. Each stimulus and associated task cue were presented twice during the encoding session.

In the test session, all 144 objects were presented again, either with the same task cue as during encoding or with a different task cue, creating two task switch conditions in which either the classification task (size vs. mechanical properties) or the response side for that stimulus changed. All encoded stimuli were randomly assigned to one of three conditions (repeat/action switch/classification switch). Participants were instructed to respond as quickly and accurately as possible throughout the task. Following this test of S-R memory, explicit recognition memory for the objects and their associated task cues was tested (Fig. 1A). Object recognition testing comprised the presentation of the 144 objects of the S-R task alongside 144 new objects in random order, and for each object the participant was asked to decide whether it was old or new. Each object was displayed for a maximum of 5 s or until a response was given. The ‘A’ and ‘L’ keys on a keyboard were used to indicate ‘new’ and ‘old’, respectively. After each decision, participants rated the confidence in their decision by pressing number keys on a standard keyboard, ranging from 1 (very confident) to 4 (guess). At the end of a trial, participants received feedback (‘correct’, ‘wrong’) for 500 ms. They were instructed to respond as accurately as possible on the task.

In order to test recognition memory of the task cue associated with the presentation of an object during the encoding session, each of the 144 objects was presented together with two task cues in random order. The task cues were positioned to the left and right of the object. Importantly, one of the task cues (e.g., S/B) had been shown together with the object during encoding, whereas the other (e.g., M/N) had not been shown. Participants were asked to indicate by a left/right key press which task cue had been associated with the object during the encoding session. Each picture was displayed until a response was made, and participants were again asked to rate the confidence in their decision as in the object recognition test. They were instructed to respond as accurately as possible during the task and did not receive feedback.

### Context manipulation

Encoding and test sessions could take place in one or two different environmental contexts. In the real environment (RE) context, participants were seated ∼50 cm away from a computer monitor while performing the experimental tasks. In the virtual reality (VR) context, participants performed the task in a virtual 3D space. The 3D space was developed using Unreal Engine 5 (version 5.1, Epic Games, USA) and displayed through VR headsets (Varjo Aero, Varjo GmbH). In this VR setting, participants sat at a table wearing VR headsets and entered a virtual space consisting of an expansive desert under a clear sky. They could move freely within the desert using a controller. In the center of the arena was a virtual table with a computer screen on it. The S-R task programs were screen-captured and mirrored onto the virtual screen via the OWL Live-Streaming Toolkit (Off World Live Limited). Thus, participants performed the same tasks as in the RE context by viewing the virtual screen and responding using a standard keyboard. During task performance, their position was fixed in front of the virtual screen, but they could still freely turn their heads. The relative distance between participants and the virtual screen was adjusted to match that of the RE context. VR and RE contexts were additionally associated with different experimenters and different labs which were approximately two kilometers apart. Allocation of the two contexts and their order to the conditions and sessions was counterbalanced across participants.

### Analysis of memory performance

Analysis of reaction times (RTs) on the S-R task during the test session followed standard procedures as described previously (Miao et al., 2023). Analyses of RTs were restricted to well-learned trials, i.e., stimuli that were correctly answered on both encoding trials, and also correctly answered in the test session. Data deviating more than two standard deviations (SDs) from the respective group mean were identified as outliers. To avoid potential influences from fast guesses or lack of attention, the 5% of trials above or below two SDs from the individual mean RT for the respective task condition (repeat/action switch/classification switch) were also discarded (Berger & Kiefer, 2021). S-R memory strength was assessed in terms of switch costs, i.e., the slowing of responses to stimuli that were presented at the test session with a task cue different from that during the encoding session, in comparison with RTs to stimuli presented with the same task cue at the test and encoding sessions. Better memory for the encoded association is thus reflected by an increase in switch costs. Switch costs were calculated across switch conditions as: 0.5 * *RT*_*action* *switch*_ + 0.5 * *RT*_*classification* *switch*_ - *RT*_*repeat*_. Analysis of performance accuracy on the S-R memory test was based on all trials (including errors and omissions).

For analyzing recognition performance for objects and task-cues, data fell above or below two SDs from the respective group mean were identified as outliers. To minimize possible confounding influences of re-encoding of the stimuli during prior S-R memory testing, we again restricted these analyses to trials that were correctly answered twice during encoding (as for the analysis of S-R memory). Moreover, these analyses were restricted to switch trials.

### Sleep recordings

Post-encoding sleep in the Sleep groups was monitored using standard polysomnography including electroencephalographic (EEG) recordings from frontal (F3, Fz, F4), central (C3, Cz, C4) and parietal (P3, Pz, P4) electrode sites according to the international 10-20 system. Additionally, the diagonal electrooculogram (EOG) and the submental electromyogram (EMG) were recorded. A ground electrode was attached to the nasion. All electrodes were referenced to linked electrodes attached to the mastoids (A1, A2). Signals were digitized at a sampling rate of 500 Hz with a BrainAmp DC system (Brain Products GmbH, Germany), and filtered using fourth-order zero-phase-shift Butterworth filters as implemented in BrainVision Analyzer 2.1). The EEG and EOG were filtered between 0.16 and 35 Hz, and the EMG between 0.16 and 70 Hz. In addition, a 50-Hz notch filter was applied to all signals.

Visual sleep scoring was conducted offline supported by custom Matlab scripts and performed based on EEG (C3 and C4) EOG, and EMG data in 30 s epochs, according to standard criteria (Rechtschaffen & Kales, 1968). Sleep EEG measures were extracted using the YASA toolbox (version 0.6.5) implemented in python (Vallat & Walker, 2021). Epochs with movement artifacts were visually identified and excluded from further analyses. Two channels from one participant were rejected due to poor recording quality. Total sleep time (TST), movement time (MT), latencies of S2 and SWS sleep, and the time in different sleep stages were determined for each participant and summarized for each context condition in Table 1. In addition, sleep spindles and slow oscillations (SOs) were summarized for each context condition in Table 2.

**Table 1.** Sleep architecture. Mean (SEMs in parentheses) duration (in min) of sleep and sleep stages. TST - Total sleep time, MT - Movement time, NonREM sleep stages S1, S2, slow wave sleep (SWS). Latency (in min) - first occurrence of sleep stage with reference to sleep onset.

| Sleep parameters | Experimental groups |  |
| --- | --- | --- |
|  | Same-Context | Different-Context |
| TST | 112.8 (2.6) | 108.5 (3.6) |
| MT | 3.8 (1.5) | 8.2 (3.6) |
| S2 latency | 9.5 (0.8) | 13.0 (1.7) |
| SWS latency | 34.0 (5.9) | 32.3 (4.5) |
| S1 | 23.5 (3.4) | 21.2 (3.0) |
| S2 | 53.4 (3.5) | 54.2 (4.6) |
| SWS | 24.7 (3.7) | 25.0 (4.3) |
| REM | 11.2 (2.1) | 8.1 (1.9) |

**Table 2.** Mean (SEMs in parentheses) densities and amplitudes of sleep spindles and slow oscillations (SOs) for participants tested in the same or different context as during encoding, averaged across the frontal, central, and parietal recording sites.

| Event type | Parameter | Location | Experimental groups |  |
| --- | --- | --- | --- | --- |
|  |  |  | Same-Context | Different-Context |
| Spindles | Density | Frontal | 1.38 (0.20) | 1.51 (0.26) |
|  |  | Central | 1.39 (0.23) | 1.58 (0.26) |
|  |  | Parietal | 1.45 (0.24) | 1.54 (0.26) |
|  | Amplitude | Frontal | 70.52 (1.81) | 69.84 (2.89) |
|  |  | Central | 68.53 (2.02) | 72.53 (2.97) |
|  |  | Parietal | 64.74 (2.02) | 70.66 (2.80) |
| SOs | Density | Frontal | 8.90 (1.47) | 8.50 (1.42) |
|  |  | Central | 7.27 (1.21) | 7.29 (1.15) |
|  |  | Parietal | 6.74 (1.22) | 6.77 (1.10) |
|  | Amplitude | Frontal | 127.64 (2.12) | 131.77 (2.31) |
|  |  | Central | 126.12 (2.29) | 131.42 (2.40) |
|  |  | Parietal | 121.65 (2.54) | 129.31 (2.38) |

Specifically, spindles were automatically identified during NonREM sleep using the EEG broadband signal if all of three thresholds were met: First, the relative power (sigma band, 11-16 Hz/ broadband, 1-30 Hz) exceeded 20%; second, the moving correlation between sigma band and broadband signals exceeded 0.65; and third, the moving root mean square (RMS) of the sigma-filtered signal exceeded the mean by 1.5 SD. Candidate spindle events identified in this manner were designated as actual spindles if their duration was between 0.5 s and 2 s, and spindle events occurring <0.25 seconds apart were merged. Absolute spindle counts were divided by the time spent in NonREM sleep to obtain spindle density per minute, and spindle amplitude was determined as the maximum difference between negative and positive voltage during each spindle event.

Moreover, as implemented in YASA (Vallat & Walker, 2021), SOs were automatically identified during NonREM sleep based on previous works (Carrier et al., 2011; Massimini et al., 2004). Specifically, a bandpass-filter from 0.3-1.5 Hz was applied to all S2 and SWS epochs. Zero-crossings in the filtered signal were determined and negative peaks with amplitudes between −40 and −200 µV, followed by positive peaks with amplitudes between +10 and +150 µV were marked as candidate events. They were retained as true SOs if all of the following conditions were met: (i) negative half-wave duration between 0.3 and 1.5 s; (ii) positive half-wave duration between 0.1 and 1 s; (iii) peak-to-peak amplitude between 75 and 350 µV. For the retained events, the zero-crossing preceding the negative peak was defined as its onset. Absolute SO counts were divided by time spent in NonREM sleep to obtain SO density per minute. Spindle-SO coupling events were identified if the center peak of the spindle fell in a ± 1.2 s window around the negative peak of the SO.

### Statistical analysis

Statistical analyses were conducted using R (version 4.4.0, R Core Team, 2021). The significance level of all statistical tests was set at 5%, and two-tailed results were used unless noted otherwise. We used a linear mixed model and a generalized linear mixed model to analyze RT and accuracy data, respectively (Baayen et al., 2008; Bates et al., 2015). The lmerTest package was used to obtain significance values via Satterthwaite’s approximation of degrees of freedom (Kuznetsova et al., 2017). Both models were fitted using restricted maximum likelihood (REML) and maximum likelihood (Laplace approximation) methods, respectively.

For the analysis of S-R memory (switch costs), in the main experiment, the model design comprised task switch (repeat/action switch/classification switch) as within-subjects factor and two between-subjects factors: group (Sleep/Wake) and context (Same/Different). We ran the model with these three factors and their interactions as fixed effects, and a random effect structure consisting of by-subject random intercepts, by-item random intercepts, and by-item random slopes for the task type (mechanical/non-mechanical vs. smaller/bigger). In the analyses of the supplementary experiments, the model did not include the group factor but still comprised task switch (repeat/action switch/classification switch) as within-subjects factor and context (Same/Different) as between-subjects factor. For the analysis of the temporal dynamics of wake-dependent enhancements in S-R memory, the model comprised task switch (repeat/action switch/classification switch) as within-subjects factor and group (Sleep/Wake/Immediate-Wake/Delayed-Sleep) as between-subjects factor. We ran the model with these two factors and their interaction as fixed effects, and the same random effects structure as above. These models account for potential variability in baseline levels and factor-level-specific evolution of performance across participants and items.

Planned contrasts were used to assess whether switch costs were significantly different from zero for the different groups; whether the effect of context varied depending on sleep and wakefulness; and whether switch costs differed between groups. Performance accuracy during testing was analyzed with a model including the same fixed and random effects, using a binomial link function to account for the binomial distribution of the categorical dependent variable.

Recognition of objects and associated task cues was similarly analyzed using a generalized linear mixed model with a binomial link function for each test. Two between-subjects factors, group and context, and their interaction were included as fixed effects, with by-item random intercepts as random effects. Other random effects were removed due to singular-fit issues, suggesting that most of the variance was captured by the fixed effects. Planned contrasts were used to investigate the same questions as for the S-R test.

Differences in sleep parameters (macro-architecture) between the Sleep groups were analyzed using independent-samples t-tests. Differences in spindle and SO parameters (density, amplitude) between the Sleep groups were analyzed using three-way mixed ANOVA, with context (Same/Different) as between-subjects factor and sleep stage (S2/SWS) and electrode site (Frontal/Central/Parietal) as within-subjects factors. To relate S-R switch costs to sleep spindles, SOs and SO-spindle couplings during post-encoding sleep, we computed Pearson correlations. Medians across the three frontal, central and parietal electrode sites, respectively, were used. Multiple comparison correction was applied for each measure with significant results, based on six correlations (calculated across the three electrode sites and two context conditions) using the Benjamini-Hochberg procedure (Benjamini & Hochberg, 1995). Correlation coefficients were compared after applying Fisher’s Z transformation as implemented in the R package cocor (Diedenhofen & Musch, 2015).

## Results

To assess context-dependency, four groups of participants performed encoding and testing on the S-R task in either the same or different contexts (Figure 1C). The context manipulation involved (i) different laboratories located in different buildings at different university campuses; (ii) different experimenters at the different locations; and (iii) different display settings, with one lab equipped with standard computer hardware and the other with a VR setup. Allocation to the four possible conditions of RE contexts vs. VR contexts during encoding vs. testing was random and counterbalanced.

### Sleep enhances S-R memory tested in the same context, wake enhances S-R memory tested in a different context

We analyzed memory of S-R associations in terms of reaction time (RT) “switch costs” at testing, i.e., the slowing of RTs in response to an object presented with a different task cue than during training (switches), in comparison to RTs for objects presented with the same task cue as during training (repeats). Thus, high switch costs indicate strong memory for the originally learned S-R association that interfere with current task performance. We collapsed action-switch and classification-switch costs, as there were no significant differences between these conditions (*b* = −17.17, *t* = −0.97, *p* = 0.333). A prior analysis confirmed that RTs to repeats did not differ between the groups (*F*(3,59) = 1.06, *p* = 0.375, Fig. S1A), excluding that the switch cost-measure was confounded by baseline differences. Also, S-R performance during the encoding session did not reveal any significant differences between groups (*F*(3,60) = 1.11, *p* = 0.351, Fig. S2A).

The effect of the retrieval context on memory differed between Sleep and Wake groups. In comparison with testing in the same context as during encoding, testing in a new context produced an increase in switch costs after the wake retention interval, indicating enhanced memory for the previously encoded S-R associations. In contrast, after the sleep retention interval, switch costs were lower in the new context, indicative of a reduced strength of the originally encoded S-R associations (*b* = −50.09, *t* = −3.29, *p* = 0.001, for interaction contrast in linear mixed model analysis; see Figure 2A for simple contrasts between groups).

**Figure 2.**
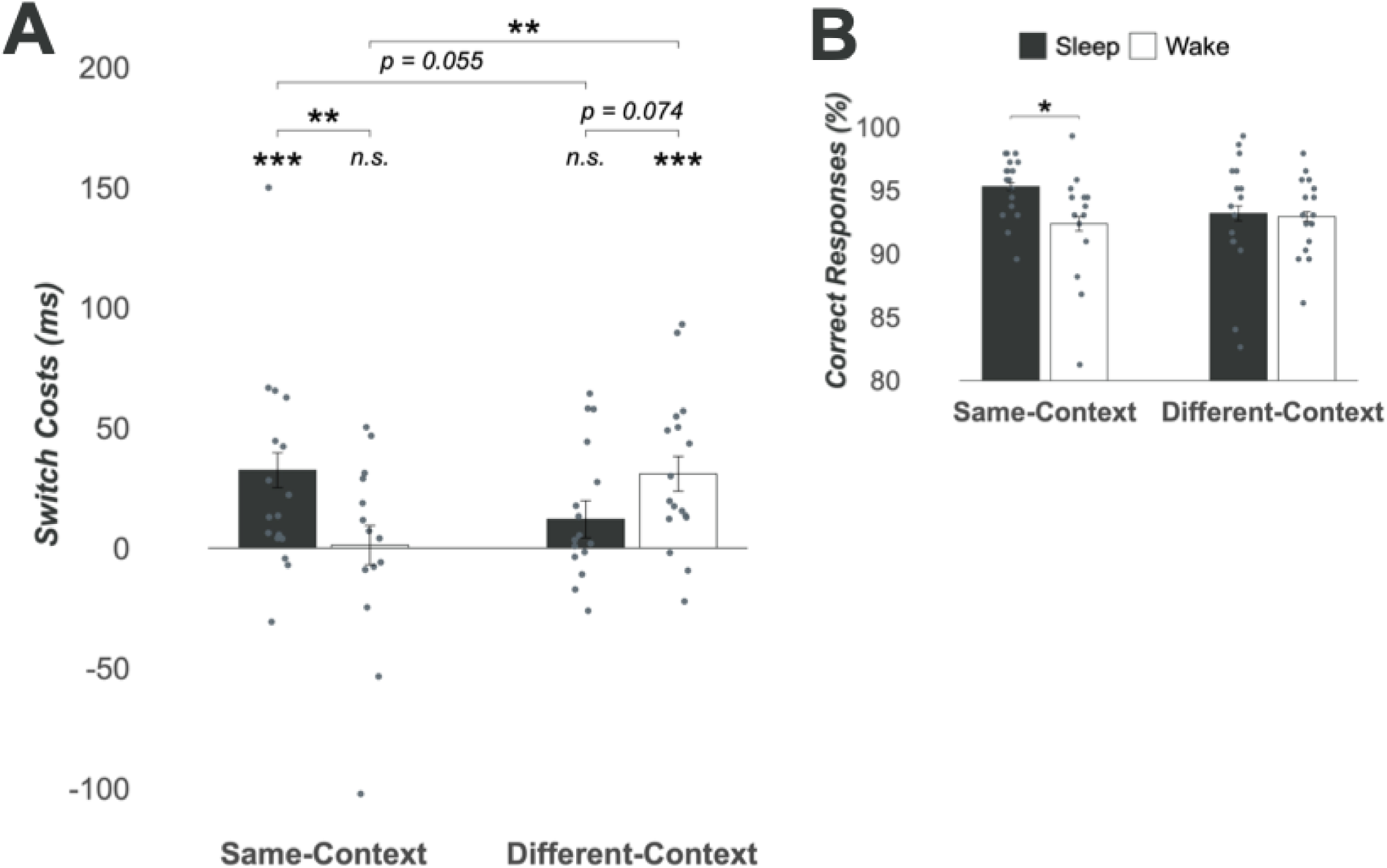
S-R memory tested in the encoding context or a different context for Wake (white) and Sleep (black) groups. **(A)** Mean ± SEM RT switch costs as indicator of memory strength are shown with individual dots overlaid. Asterisks above individual bars indicate significant difference from zero, those above brackets indicate significance for simple contrasts. **(B)** Accuracy of performance (% correct responses) shown in the same way with individual dots overlaid. ***, P < .001; **, P < .01; *, P < .05.

This pattern was confirmed in more focused comparisons: Examining the groups tested in the original context revealed higher switch costs after sleep than wakefulness (*b* = 31.17, *t* = 2.85, *p* = 0.004). This was due to the presence of significant switch costs after sleep (contrast against zero, *b* = 32.40, *t* = 4.45, *p* < 0.001), whereas switch costs after wake did not differ from zero (*b* = 1.23, *t* = 0.15, *p* = 0.880). Conversely, examining the groups tested in a new context revealed a trend for higher switch costs following wakefulness (*b* = −18.92, *t* = - 1.78, *p* = 0.074), with significant switch costs only in the Wake group (*b* = 30.91, *t* = 4.26, *p* < 0.001), but not in the Sleep group (*b* = 11.99, *t* = 1.55, *p* = 0.121).

Comparisons between the Sleep groups indicated a trend towards greater switch costs when the context was the same during encoding and testing (*b* = −20.41, *t* = −1.92, *p* = 0.055). Conversely, in the Wake groups, switch costs were significantly greater when the context changed from encoding to testing (*b* = 29.68, *t* = 2.71, *p* = 0.007). Notably, switch costs in the Wake group tested in the different context were comparable to those of the Sleep group tested in the original context (*b* = −1.50, *t* = −0.15, *p* = 0.884).

In terms of performance accuracy, the Sleep group displayed better performance than the Wake group when the context remained constant (*b* = 0.53, *z* = 2.44, *p* = 0.015, Fig. 2B). However, there was no group difference when the context changed between encoding and testing (*b* = 0.10, *z* = 0.49, *p* = 0.624). Given the differences between Sleep and Wake groups in accuracy, in additional linear mixed model analyses, we assessed a potential confounding influence of speed-accuracy trade-offs on RT switch-costs, incorporating the accuracy for trials with a changed task cue as a covariate. All results reported above remained unchanged, in line with the view that participants did not sacrifice response speed for accuracy.

### The wake-dependent enhancement in S-R memory tested in a different context has a fast onset and is reversed by sleep

Given the surprising finding of post-encoding wakefulness enhancing S-R memory when tested in a different context, we explored the temporal dynamics of this effect in two supplementary experiments. In the first of these, participants were tested in the same or different context closely after (∼20 min) encoding (Immediate-Wake groups). Significant switch costs were observed only in the group tested in a different context (*b* = 31.49, *t* = 4.91, *p* < 0.001), but not in the group tested in the same context (*b* = 5.11, *t* = 0.78, *p* = 0.434), and switch costs significantly differed between groups (*b* = 26.38, *t* = 2.88, *p* = 0.004; Fig. S3A). Performance accuracy (Fig. S3B) as well as control measures (RTs to repeats, Fig. S1B; S-R performance during encoding, Fig. S2B) were comparable between contexts (all *p* > 0.314).

The second supplementary experiment examined the persistence of the wake-dependent increase in S-R memory. Participants remained awake for 12 hours after encoding and S-R memory was tested in a different context on the next day, after nocturnal sleep (Delayed-Sleep). This test took place 24 hours after encoding and revealed no significant switch costs (*b* = 5.09, *t* = 0.75, *p* = 0.453, Fig. 3B). Indeed, the Delayed-Sleep group showed switch costs comparable to those seen in Sleep participants in the main experiment when tested in a different context (*b* = −6.08, *t* = −0.62, *p* = 0.534), and performance accuracy was even higher in the Delayed-Sleep group than in the main-experiment Sleep group (*b* = 0.54, *z* = 2.24, *p* = 0.025). This indicates that sleep, even when occurring 12 hours after encoding, reverses the enhancing effects of wakefulness on S-R memory tested in a different context.

**Figure 3.**
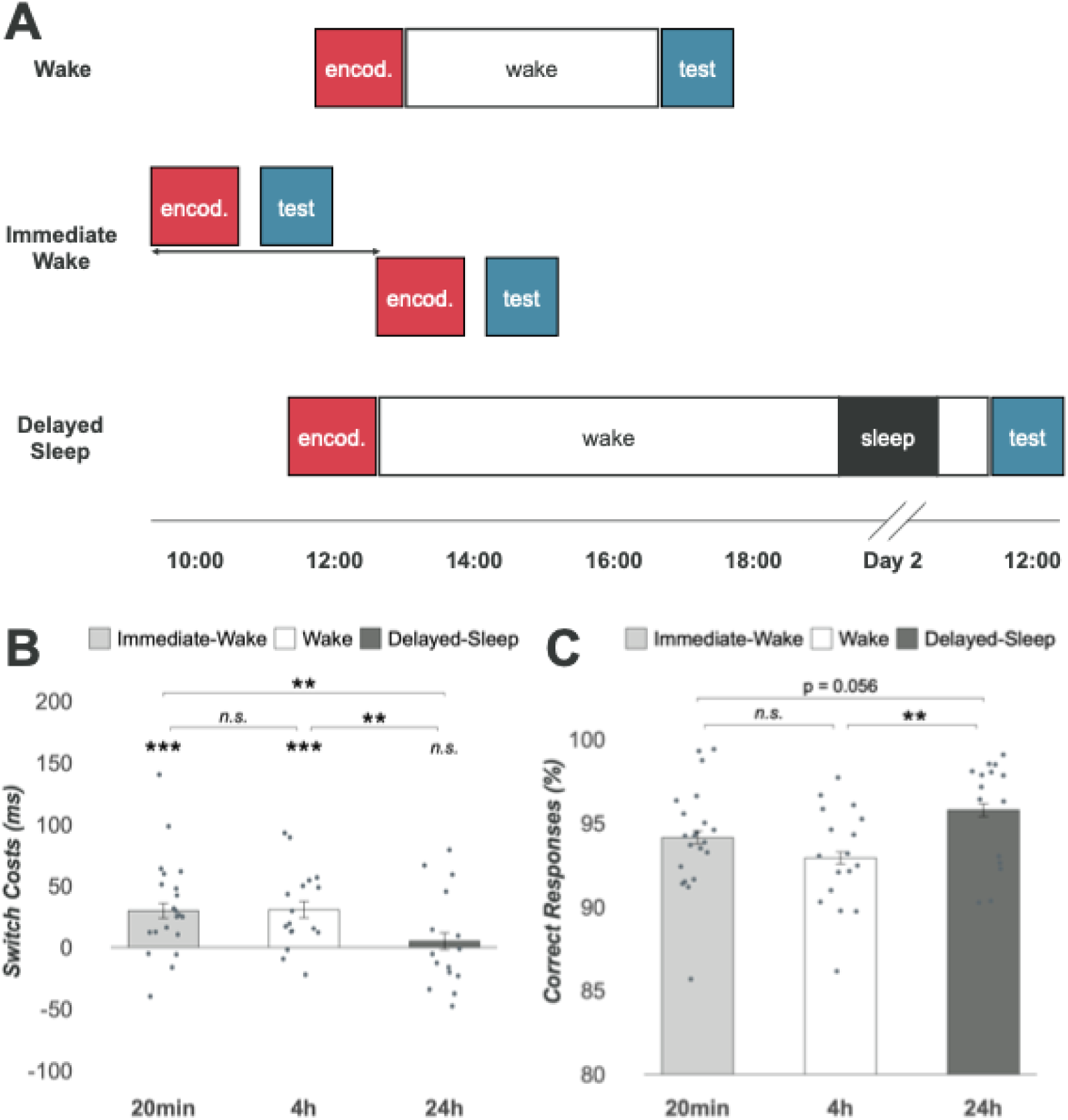
S-R memory tested in in a different context across main and supplementary experiments. **(A)** Timelines for the Wake group (main experiment) and Immediate-Wake and Delayed Sleep groups (supplementary experiments). Different colors indicate context difference. In the Immediate-Wake group, encoding started between 10:00 and 13:00, took approximately 1h, and was followed by testing after 20 min. **(B)** Mean ± SEM RT switch costs are shown with individual dots overlaid. Asterisks above individual bars indicate significant difference from zero, those above brackets indicate significance for simple contrasts. **(C)** Accuracy of performance (% correct responses) shown in the same way with individual dots overlaid. Endod., encoding; ***, P < .001; **, P < .01; *, P < .05.

A comparison of switch costs in the new context across experiments (i.e., Wake, Immediate-Wake, and Delayed-Sleep groups) confirmed that the wake-dependent increase in S-R memory has a fast onset but is reversed by delayed sleep (Fig. 3B). Specifically, switch costs were similar between Immediate-Wake and Wake groups (*b* = 0.90, *t* = 0.10, *p* = 0.919), but significantly reduced in the Delayed-Sleep group in comparison to both the Immediate-Wake (*b* = 25.03, *t* = 2.77, *p* = 0.006) and Wake groups (*b* = 25.93, *t* = 2.73, *p* = 0.006). Similarly, performance accuracy did not reveal differences between the Immediate-Wake and Wake groups (*b* = −0.22, *z* = −1.08, *p* = 0.279). In contrast, it was higher in the Delayed-Sleep group compared to the Wake group (*b* = 0.66, *z* = 2.80, *p* = 0.005), and showed a trend toward being higher than in the Immediate-Wake group (*b* = 0.44, *z* = 1.91, *p* = 0.056; Fig. 3C).

### Sleep binds memory for objects and task cues to the encoding context

After the S-R memory test, participants in the main experiment performed two recognition tests (Fig. 4). In the first, they were presented with images of the original objects interspersed with an equal number of novel objects, and asked to classify each object as old or new. In the second recognition task, participants were presented with the learned objects together with two sets of task cues and were asked which of the cue displays had been associated with the object during the encoding phase. To minimize confounding effects of re-encoding during the preceding tests, analyses of recognition performance were restricted to stimuli for which correct responses had been given during the encoding session, indicating effective encoding of the original memory trace, and that were presented as switch trials during the S-R memory test. (Corresponding results for the Immediate-Wake and Delayed-Sleep groups of the supplementary experiments are given in Fig. S4).

**Figure 4.**
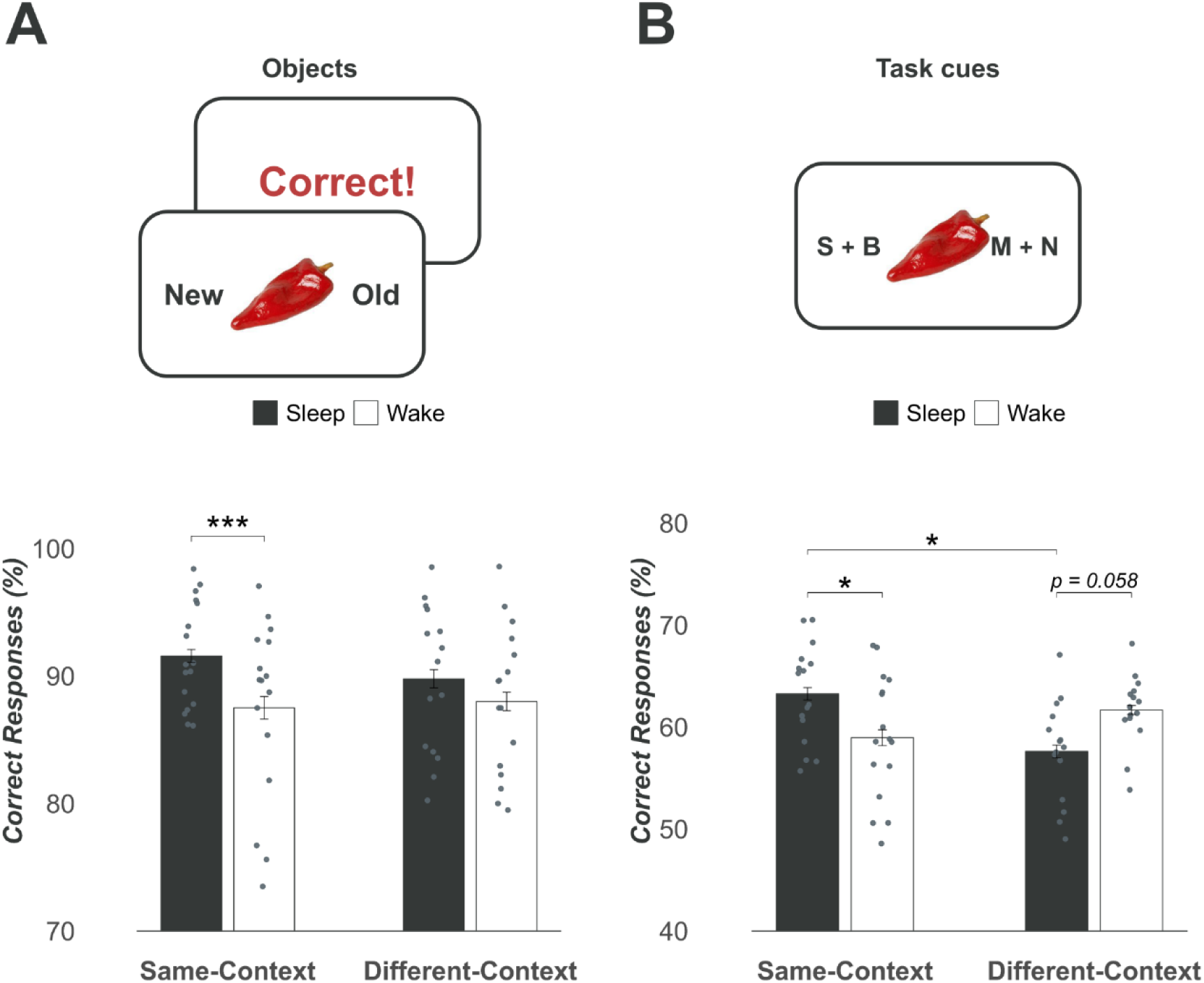
Recognition memory for objects **(A)** and task cues (**B)** tested in the encoding context or a different context for Wake (white) and Sleep (black) groups. Participants responded by pressing buttons on either the left or right side. They were required to make a confidence judgment after each response. Feedback was given after each trial on the object recognition task. Mean ± SEM correct response (%) is shown with individual dots overlaid. Comparisons are simple contrasts. ***, P < .001; **, P < .01; *, P < .05.

Object recognition was better in the Sleep than Wake group (*b* = 0.32, *z* = 3.36, *p* < 0.001). This difference was not affected by the testing context (interaction contrast, *b* = −0.27, *z* = −1.40, *p* = 0.162). Given the numerical differences between context conditions, we nevertheless calculated separate contrasts which revealed a significant difference when the context remained the same during encoding and testing (*b* = 0.46, *z* = 3.39, *p* < 0.001, Fig. 4A), whereas both groups demonstrated comparable performance when the context was changed (*b* = 0.19, *z* = 1.37, *p* = 0.170).

Recognition of the task cue associated with an object during encoding differed between Sleep and Wake groups depending on the testing context (interaction contrast, *b* = −0.34, *z* = −2.83, *p* = 0.005, Fig. 4B). Like object recognition, recognition of the associated task cue was better in the Sleep than Wake group when the test context was the same as during encoding (*b* = 0.18, *z* = 2.11, *p* = 0.035). In the new context, this difference was reversed, but only marginally significant (*b* = −0.16, *z* = −1.90, *p* = 0.058). In addition, the Sleep group tested in the same context performed better than the Sleep group tested in the changed context (*b* = −0.22, *z* = −2.56, *p* = 0.010), whereas no such difference was seen between the wake groups (*b* = 0.12, *z* = 1.43, *p* = 0.153). Overall, this pattern indicates that consolidation during sleep enhances recognition performance over wake consolidation, particularly when the context at testing remains the same as during encoding.

### NonREM sleep spindles predict S-R memory in the same context

Additional analyses aimed to relate improved S-R memory (i.e., enhanced switch costs) in the Sleep group tested in the same context as during encoding to the architecture of post-encoding sleep. The two Sleep groups did not differ in the macro-architecture of sleep (Table 1, all *p* > 0.070), and there were no differences between groups in NonREM sleep spindle and slow oscillation (SO) densities, i.e., EEG oscillatory events commonly associated with memory processing (Table 2, for main effect and related interactions, all *p* > 0.383).

Spindle density was positively correlated with switch costs when participants were tested in the original encoding context (frontal, *r* = 0.63, *t*(15) = 3.13, *p* = 0.007; central, *r* = 0.50, *t*(15) = 2.23, *p* = 0.041; parietal, *r* = 0.54, *t*(15) = 2.46, *p* = 0.027), with the correlation for frontal spindles surviving multiple comparison correction (*p* = 0.041, see Methods). In contrast, there were no significant correlations between spindle density and switch costs when testing occurred in a different context (all *p* > 0.591). Importantly, the correlation coefficients differed significantly between groups tested in the same and different context for frontal (Fisher’s *Z* = 2.14, *p* = 0.033, Fig. 5) and central spindle density (Fisher’s *Z* = 2.03, *p* = 0.042). Analyses of spindle amplitude (all p > 0.745), densities of SOs (all *p* > 0.209) and spindle-SO couplings (all *p* > 0.394) did not reveal consistent correlations with switch costs.

**Figure 5.**
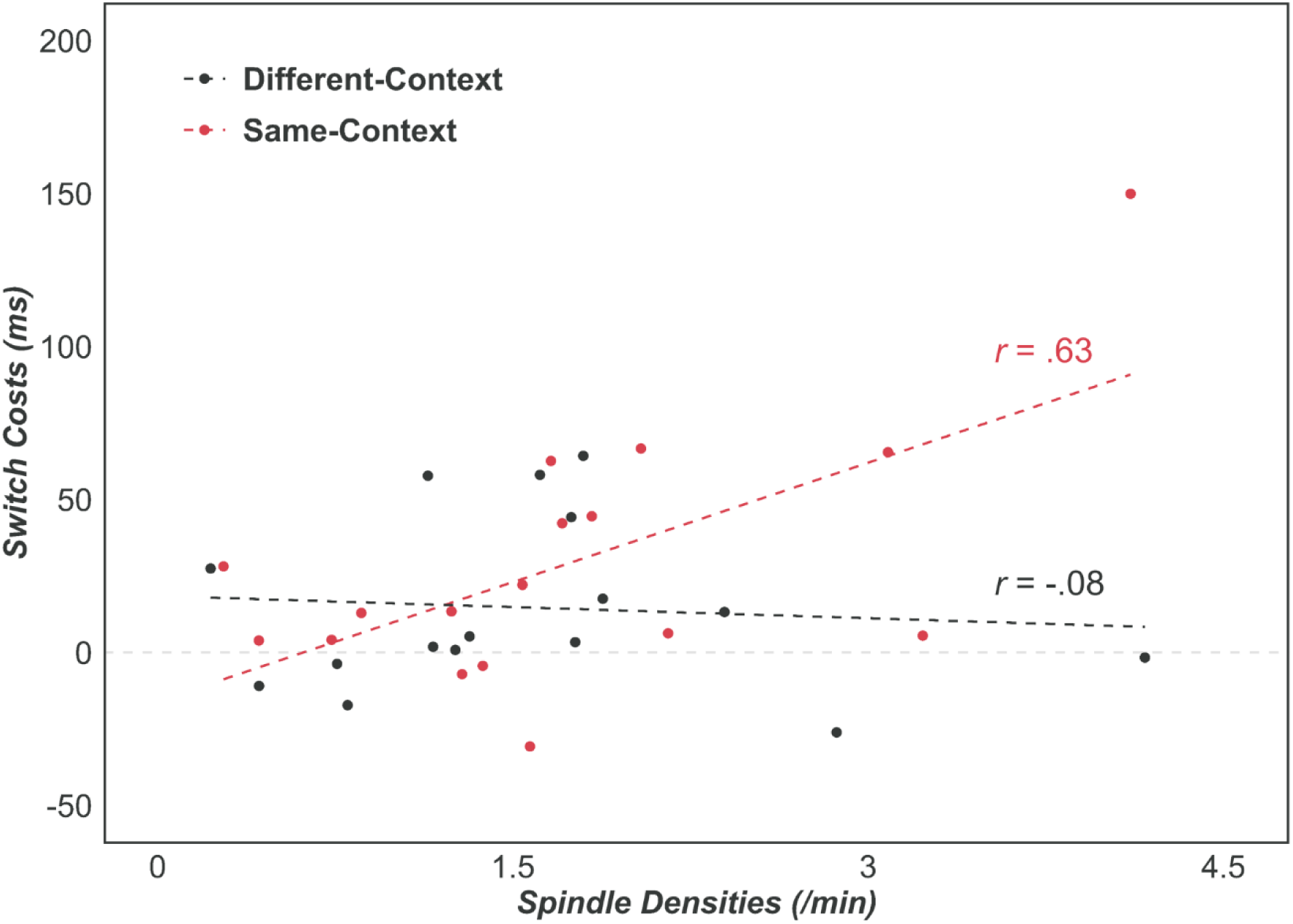
Spindle density during post-encoding NonREM sleep was positive correlated with switch costs when S-R memory was tested in the same (red) context as during encoding but not when tested in a different (black) context (Fisher’s Z = 2.14, p = 0.033, for difference between correlation coefficients). Results at the frontal recording sites are shown.

## Discussion

We assessed memory for learned stimulus-response (S-R) associations after consolidation periods of sleep and wakefulness, using reaction time (RT) switch costs as an indicator for the strength of S-R memories. Our findings indicate a dependency of consolidation effects on the context of retrieval testing: When participants slept after learning, memory for S-R associations was stronger when tested in the original learning context than in a different test context. In contrast, when participants remained awake after learning, S-R memory was stronger when retrieval was tested in a different context. Sleep and wake-associated benefits in S-R memory at retrieval in same and different contexts were of comparable size. The wake-associated benefit was already present shortly after encoding and was reversed by sleep, even if it occurred after a prolonged 12-hour wake period following encoding. Sleep spindles over the frontal cortex predicted sleep-dependent S-R memory improvements when retrieval was performed in the same context, but not in a different context. The findings point to a distinct offline-consolidation process in the wake state that maintains memory for an experienced event in a non-integrated format and, thus, differs from sleep-dependent consolidation of representations that integrate the event with its specific context in which it occurred.

The sleep-dependent benefit of S-R memory tested in the original learning context replicates our previous findings indicating that S-R associations undergo consolidation during sleep (Miao et al., 2023). In line with the present findings, this previous study also found NonREM sleep spindle activity to be positively correlated with S-R memory strength. Extending our previous findings, here we show that sleep does not benefit S-R memory when tested in a context different from learning.

Beneficial effects of sleep on recognition memory for the objects used in the S-R task, as well as for the cues associated with the objects during learning, were also largely restricted to memory testing in the original learning context. This is in line with the notion of a systems consolidation process during sleep that strengthens the binding of experienced events and items into their spatio-temporal context (Berres et al., 2024; Contreras et al., 2024; Inostroza & Born, 2013; van der Helm et al., 2011). On the other hand, these findings contradict the view of sleep de-contextualizing newly encoded episodic memories (Jurewicz et al., 2016; Lewis & Durrant, 2011). Indeed, the contextual binding of experienced events is commonly thought of as a central function of the hippocampus (Jezek et al., 2011; Kelemen & Fenton, 2010), and there is converging evidence from human and rodent studies that consolidation during sleep is driven by the neuronal replay of representations in hippocampal circuitry during NonREM sleep (Brodt et al., 2023; Gridchyn et al., 2020; Sawangjit et al., 2018; Schapiro et al., 2019). This contextual replay of hippocampal representations co-occurs with NonREM sleep spindles originating from the thalamus as a mechanism that may foster synaptic plasticity underlying the enhancing effect of sleep on item-context binding in episodic memory (Antony et al., 2019; Petzka et al., 2022).

The central finding of this study is that S-R memory after a post-encoding wake period was superior to that after post-encoding sleep when testing took place in a context different from learning. This replicates recent findings in rats which showed enhanced novel object recognition after a post-encoding wake period compared to post-encoding sleep when the rats had to recognize the test objects in a context different from that during encoding (Sawangjit et al., 2022). Those experiments also revealed that, unlike consolidation during sleep, consolidation associated with post-encoding wakefulness was enhanced, rather than impaired, by experimentally inactivating the rats’ hippocampus. This implies that under normal conditions, ongoing hippocampal activity interferes with consolidation during wakefulness.

Against this backdrop, our result of superior S-R memory after wake when tested in a novel context may be partly explained in the framework of contextual binding theory (Mednick et al., 2011; Yonelinas et al., 2019) which assumes that sleep enhances episodic memory mainly by protecting it from contextual interference. In this view, the wake state is hallmarked by continuous encoding of contextual information into hippocampal networks. This continuous encoding maps contextual drifts, such that items often become associated with multiple contexts (Howard & Kahana, 2002; Yonelinas et al., 2019). Hence, consolidation during wakefulness is expected to produce item representations that are only weakly associated with their encoding context. Consequently, when retrieval is tested in the original context, these representations are only weakly reactivated, in comparison with item representations consolidated during sleep, because sleep, in the absence of contextual interference, enables the formation of stable associations between items and their unique encoding context. However, such enhanced item-in-context memories are more strongly impacted by contextual interference when they are to be retrieved in a novel context, thus rendering retrieval more difficult than for memories consolidated during wakefulness.

A seemingly paradoxical finding of our study is that S-R associations after consolidation during wakefulness were even significantly stronger when retrieved in the different than when retrieved in the same context as during learning. This superior expression of a memory in a novel context after wake consolidation has been similarly observed in rats (Sawangjit et al., 2022) and is difficult to explain based on contextual binding and interference alone. It is reminiscent of memory enhancements observed after active inhibition of the hippocampus during wake consolidation, e.g., by alcohol (Parker et al., 1981; Sawangjit et al., 2022). It might be that reactivation of representations outside the hippocampus in the wake state is accompanied by an active suppression of context-related hippocampal activity (Albouy et al., 2013) that could interfere with a consolidation process that eventually serves to maintain event memories in an easily amendable format. In this scenario, the strength of extra-hippocampal reactivation and hippocampal suppression might be reciprocally linked, such that extrahippocampal reactivations of event memories that occur in a potentially highly interfering context – e.g., at retrieval in a novel context – are associated with a distinct suppression of context-related hippocampal inputs. Such a reciprocal link could indeed be a reflection of the competitive interaction between hippocampal and extra-hippocampal (e.g., striatal) memory systems (Albouy et al., 2008; Hartley & Burgess, 2005; Packard & Goodman, 2013; Poldrack & Packard, 2003), with recent neuroimaging evidence indicating that synchronous hippocampo-cortical reactivations may even promote forgetting (Tanrıverdi et al., 2023).

Such a scenario would also be consistent with the temporal dynamics we revealed for the wake-associated benefit of S-R memory when tested in a context different from learning. Rather than decaying over time, the size of this benefit was almost identical at tests shortly (∼20 min) after and ∼4 hours after encoding, suggesting a stabilizing effect of the wake period on S-R memories. However, the wake-associated memory enhancement disappeared as soon as sleep intervened, even when this sleep period occurred with a delay of 12 hours. This result contrasts with findings in rats, where the enhancing effect of a 2-hour post-encoding wake period on object recognition memories probed in a different context persisted for one week, apparently surviving multiple intervening sleep periods (Sawangjit et al., 2022). This discrepancy is difficult to explain, deserving further investigation. It might be that in humans, compared to rats, memory reactivations during sleep are more powerful in re-associating event representations with their respective hippocampal context representations (Hahn et al., 2024).

The wake-associated increase in S-R memory when tested in a context different from learning was remarkably robust and observed in different experiments. Yet, it stands in striking contrast with the classic finding that memory retrieval is better when tested in the same context as learning (Godden & Baddeley, 1975; Smith, 1979, 1982; Smith et al., 1978). However, those studies typically relied on declarative tasks with an explicit assessment of item-context associations. Instead, here, we used a reaction time-based priming measure to assess memory for S-R associations. Context interactions at retrieval likely differ between explicit measures of memory that more strongly rely on hippocampal function, and the priming-based measures used here (Chun & Phelps, 1999; Ramponi et al., 2011; Schott et al., 2005; Wang et al., 2010). Consistent with this explanation, we did not observe a clear enhancing effect of retrieval in a different context for explicit recall measures, i.e., the recall of task cue-stimulus associations or recognition of objects.

The concept of “context” is central to our study in that it dissociates sleep and wake modes of consolidation. However, there is currently no clear agreement upon what defines the context of an event (Easton et al., 2024; Rudy, 2009; Stark et al., 2018), which possibly limits the conclusions that can be drawn from our results. The finding that consolidation during sleep but not wakefulness leads to an enhanced event-context binding is incompatible with the notion that context is merely re-constructed at retrieval rather than being encoded (Easton et al., 2024). More commonly, it is assumed that context is encoded as such, and most such concepts agree in that contexts of events are seen as stable over time, moderately complex, and having limited (explicit or implicit) task relevance (Stark et al., 2018). Moreover, the hippocampus is commonly considered as a core structure binding item and context (Rudy, 2009; Yonelinas et al., 2019), although hippocampal involvement is not a sufficient criterion, as it also forms non-contextual associations between items. Here, we followed these definitions of context in that we varied practically all aspects of the external environment that persisted throughout the experimental episode but were not of immediate relevance to the S-R task, including the experimenter, the laboratory and its location in the town of Tübingen as well as the display technique used to present the task (virtual reality vs. standard screen). Nevertheless, given the crudeness of current concepts of context, it remains to be clarified whether one of these aspects of context – and its interactions with specific task requirements - is of particular relevance to our findings.

Our findings corroborate previous work showing that consolidation does not only occur during sleep but also in the wake state (Wamsley, 2019). By testing S-R memories in the same context as during encoding or in a different context, we unmask distinct features of consolidation in the two states. Memory testing in a different context indicated that wake consolidation processes specifically pertain to the maintenance of event memory not bound to a specific encoding context.

## Author contributions

X.M., J.B., and K.R. designed research; X.M., A-S.T., J.D. and A-L.W. performed research; X.M. analyzed data; and X.M., J.B., and K.R. wrote the paper.

## Funding statement

This work was supported by grants from the Deutsche Forschungsgemeinschaft (SFB 1233 “Robust Vision: Inference Principles and Neural Mechanisms,” Project no. 276693517, TP8 to KR; FOR 5434 “Information Abstraction during Sleep” to JB) and the European Research Council (ERC, AdG 883098, “SleepBalance” to JB).

## Conflict of interest disclosure

The authors declare no conflict of interest.

## Supplementary Information

**Figure S1.**
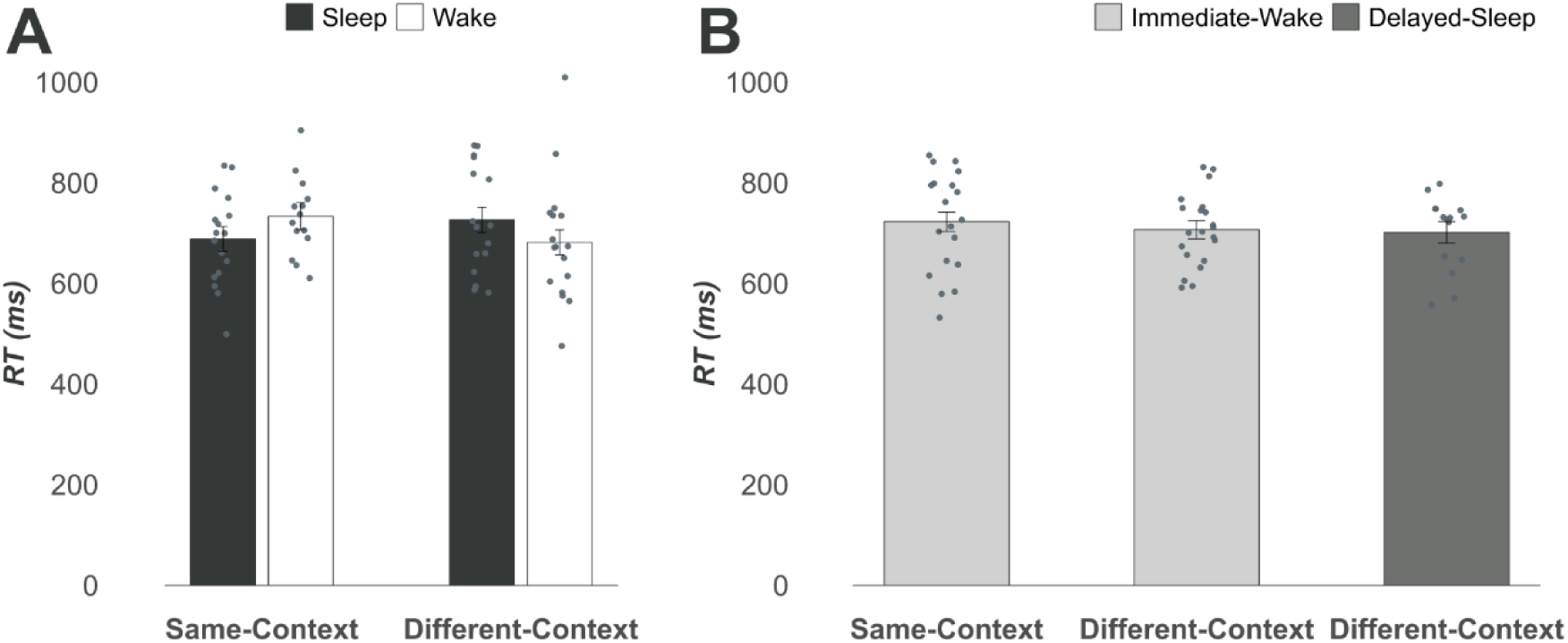
RTs of repeat trials at S-R memory test performed in the same and different contexts (A) for the Sleep and Wake groups of the main experiment, and (B) for the Immediate-Wake and Delayed-Sleep groups of the supplementary experiments. (The Delayed-Sleep group was only tested in the different context).

**Figure S2.**
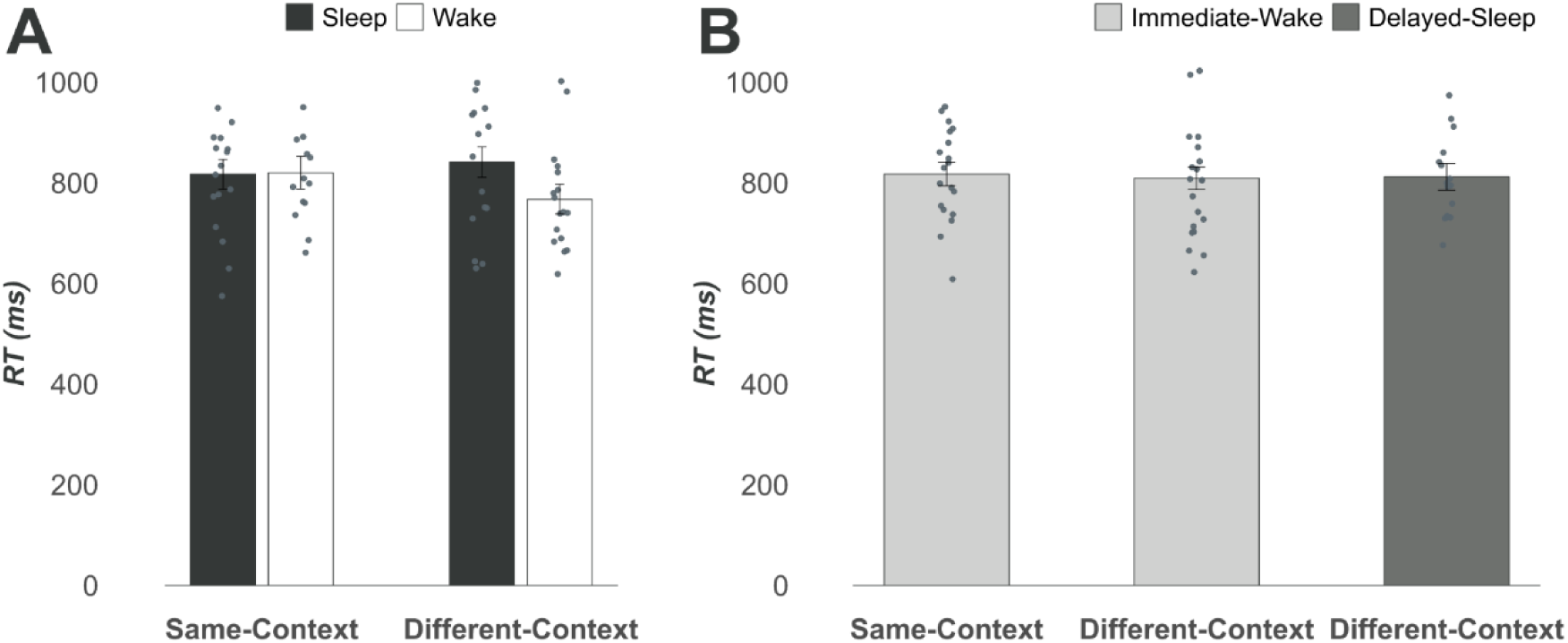
S-R memory performance at encoding, separately for the groups tested in the same or different context, (A) for the Sleep and Wake groups of the main experiment, and (B) for the Immediate-Wake and Delayed-Sleep groups of the supplementary experiments. (The Delayed-Sleep group was only tested in the different context).

**Figure S3.**
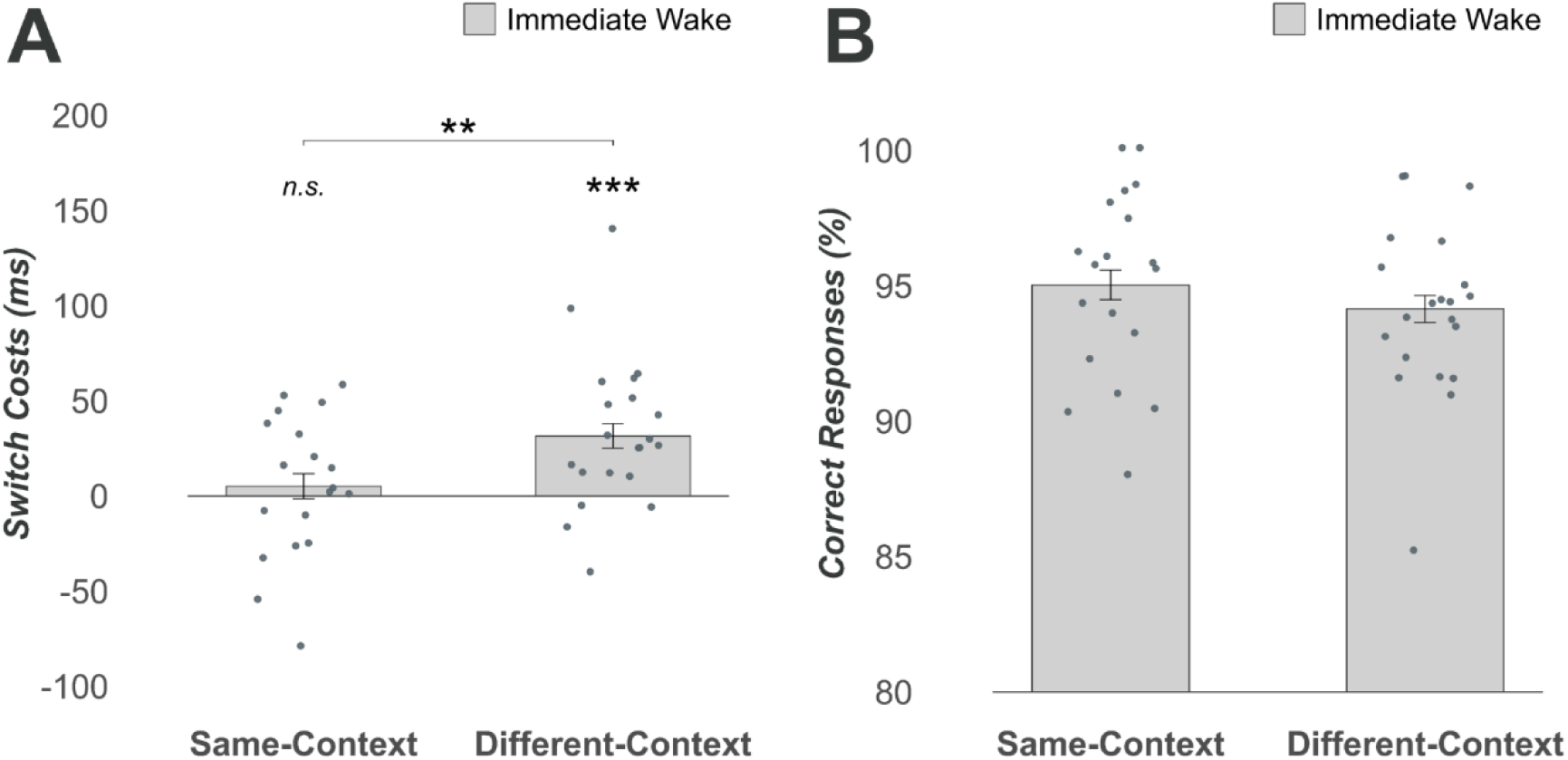
Memory performance tested immediately after encoding in the same or different context as during encoding. For S-R memory, (A) Mean ± SEM RT switch costs as indicator of memory strength and (B) accuracy of performance (% correct responses) are shown with individual dots overlaid. Asterisks above individual bars indicate significant difference from zero, those above brackets indicate significance for simple contrasts. ***, P < .001; **, P < .01; *, P < .05.

**Figure S4.**
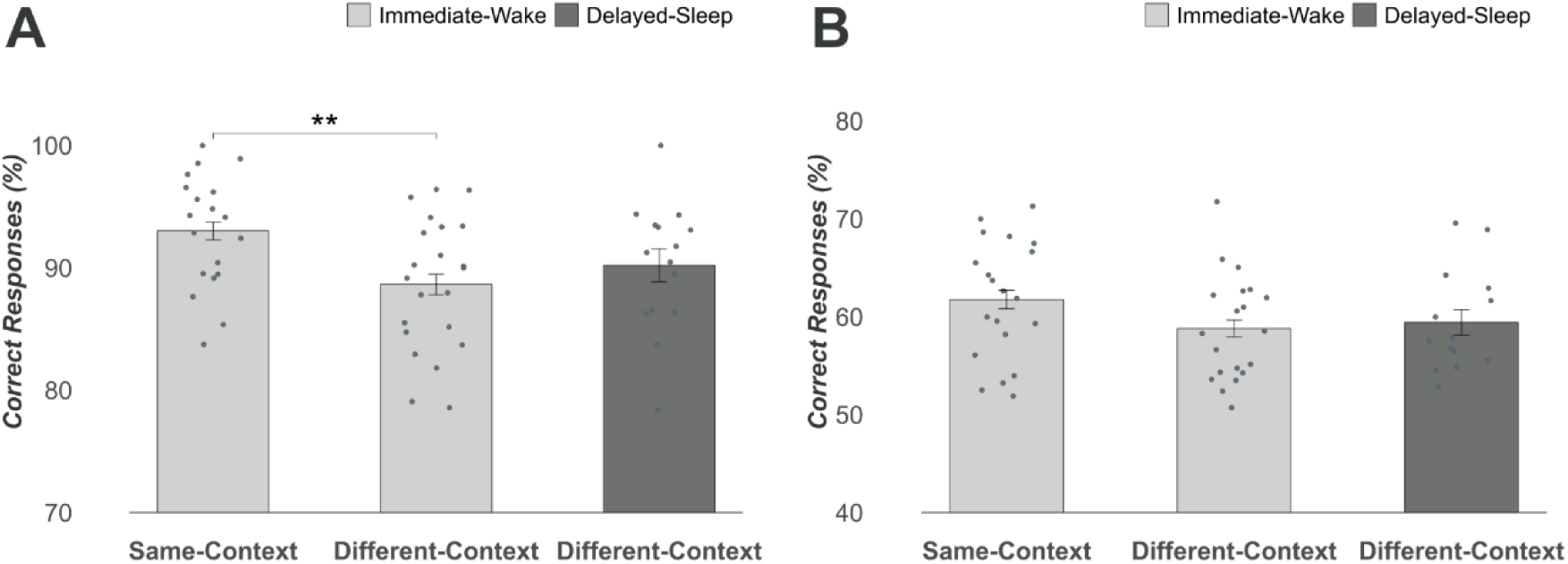
Recognition test performance. Mean ± SEM correct response (%) is shown with individual dots overlaid for (A) objects and (B) task cues for the Immediate-Wake groups tested in the same or different context and for the Delayed-Sleep group (tested in the different context. Asterisks above individual bars indicate significant difference from zero, those above brackets indicate significance for simple contrasts. ***, P < .001; **, P < .01; *, P < .05.

## Notes

### Competing Interest Statement

The authors have declared no competing interest.

